# Single-molecule studies reveal the off-pathway elemental pause state as a target of streptolydigin inhibition of RNA polymerase and its dramatic enhancement by Gre factors

**DOI:** 10.1101/2023.06.05.542125

**Authors:** Anatolii Arseniev, Mikhail Panfilov, Georgii Pobegalov, Alina Potyseva, Polina Pavlinova, Maria Yakunina, Jookyung Lee, Sergei Borukhov, Konstantin Severinov, Mikhail Khodorkovskii

**Affiliations:** Peter the Great St. Petersburg Polytechnic University, Saint Petersburg, Russia; Institute of Molecular Genetics, Russian Academy of Sciences, Moscow, Russian Federation; Department of Cell Biology and Neuroscience, Rowan University School of Osteopathic Medicine, Stratford, NJ 08084-1489, USA; Institute of Gene Biology, Russian Academy of Sciences, Moscow, Russia; Waksman Institute of Microbiology, Rutgers, The State University of New Jersey, Piscataway, NJ, United States

## Abstract

Antibiotic streptolydigin (Stl) inhibits bacterial transcription by blocking the trigger loop folding in the active center of RNA polymerase (RNAP), which is essential for catalysis. We use acoustic force spectroscopy to characterize the dynamics of transcription elongation in ternary elongation complexes of RNAP (ECs) in the presence of Stl at a single-molecule level. We found that Stl induces long-lived stochastic pauses while the instantaneous velocity of transcription between the pauses is unaffected. Stl enhances the short-lived pauses associated with an off-pathway elemental paused state of the RNAP nucleotide addition cycle. Unexpectedly, we found that transcript cleavage factors GreA and GreB, which were thought to be Stl competitors, do not alleviate the streptolydigin-induced pausing; instead, they synergistically increase transcription inhibition by Stl. This is the first known instance of a transcriptional factor enhancing antibiotic activity. We propose a structural model of the EC-Gre-Stl complex that explains the observed Stl activities and provides insight into possible cooperative action of secondary channel factors and other antibiotics binding at the Stl-pocket. These results offer a new strategy for high-throughput screening for prospective antibacterial agents.

## INTRODUCTION

DNA-dependent RNA polymerase (RNAP) is the key enzyme of the transcription process and is essential for life. In bacteria, a single multisubunit RNAP synthesizes all types of RNAs, including mRNA, tRNA, rRNA, and noncoding RNAs. During all stages of the transcription cycle, the enzyme’s function is regulated by a large variety of transcription factors and small RNAs. RNAP’s activity is also modulated by several cellular metabolites and secondary messengers (Artsimovitch et al., 2003; Gourse et al., 2018; Laptenko et al., 2003; Perederina et al., 2004; Vassylyev et al., 2007). Being at the center of gene expression, bacterial RNAP is a proven target for a wide range of antibiotics (Artsimovitch et al., 2003; Ma et al., 2016; Mosaei and Harbottle, 2019; Yuzenkova et al., 2013). Understanding the mechanisms of their action and resistance is crucial for developing new generations of RNAP inhibitors. Structural and biochemical data show that most RNAP-targeting drugs disrupt (sterically or allosterically) the enzyme’s interactions with the DNA template, the RNA transcript, the binding of NTP substrates, and/or interfere with translocation (Artsimovitch and Landick, 2002; Belogurov et al., 2009). Of these, only transcription initiation inhibitors, rifamycins and fidaxomicin/lipiarmycin, are currently approved for clinical applications (Yang et al., 2015). Several prospective drugs, including tagetitoxin, salinamide A, CBR class molecules, streptolydigin, and cyclic peptide antibiotics Microcin J25 and Capistruin, inhibit the RNAP catalytic activity (the nucleotide addition cycle, NAC), thus acting at both initiation and elongation stages of transcription. Although these inhibitors operate via distinct and complex mechanisms, ultimately, they act by obstructing the folding/movement of mobile elements of the RNAP active center essential for catalysis, including β’ bridge helix (BH), β’ F-loop (FL), β’ trigger loop (TL), and β fork-2 loop (FL2)(Artsimovitch et al., 2003; Bae et al., 2015; Malinen et al., 2014; Yuzenkova et al., 2013). Streptolydigin (Stl), produced by *Streptomyces lydicus*, is a derivative of tetramic acid and is one of the oldest known RNAP inhibitors (Crum et al., 1955; Deboer et al., 1955). It selectively targets bacterial transcription without affecting the eukaryotic RNAPs. Early studies demonstrated a high broad-spectrum activity of Stl against Gram+ bacteria (Deboer et al., 1955; Karwowski et al., 1992), making it a promising antimicrobial agent.

Stl inhibits all enzymatic activities of RNAP, including nucleotide incorporation (phosphodiester bond formation), pyrophosphorolysis, and intrinsic exo- and endonucleolytic RNA hydrolysis (Mazumder et al., 2021; McClure, 1980; Sosunov et al., 2003; Temiakov et al., 2005; Tuske et al., 2005). It also impedes DNA translocation during transcription elongation (McClure, 1980; Tuske et al., 2005) and interferes with endonucleolytic reactions catalyzed by transcript cleavage factor GreA (Temiakov et al., 2005). The crystal structures of *Thermus thermophilus* (*Tth*) RNAP (PDB: 2PPB, 1ZYR, (Temiakov et al., 2005; Tuske et al., 2005) and its elongation complex (EC) with bound Stl (Vassylyev et al., 2007) (PDB:2A6H, Vassylyev et al., 2007) revealed that the inhibitor binds through the downstream dsDNA-binding channel ∼20Å away from the catalytic center in a tight pocket formed by conserved residues of β’BH, β’TL, βFL2, and βD-loop II (**Suppl Figure S2A**). Like many other allosteric NAC inhibitors, Stl binding locks the flexible BH and TL elements in, respectively, straight and unfolded (open) conformations incompatible with catalysis (Mosaei and Harbottle, 2019; Murakami, 2015; Temiakov et al., 2005; Tuske et al., 2005; Vassylyev et al., 2007). This model is consistent with recent biophysical data (Mazumder et al., 2020), demonstrating that TL in the RNAP complex with Stl has predominantly open conformation. Unlike other known RNAP inhibitors, Stl directly contacts the nucleotides of the template and nontemplate DNA strands at the downstream edge of the transcription bubble (downstream fork junction) (Vassylyev et al., 2007)(Vassylyev 2007, **Suppl Figure S2B**). Curiously, these interactions, possibly mediated by a noncatalytic Mg^2+^-ion, contribute to Stl’s high binding affinity to *Tth* transcription complexes (Temiakov et al., 2005; Tuske et al., 2005; Zorov et al., 2014) and are presumed to be important for its inhibitory activity (Jeong et al., 2014). Although the structure of RNAP-Stl complexes is available only for the *Tth* enzyme, the substantial sequence and structural similarity it shares with *E. coli* RNAP indicates that te key residues of the Stl-binding pocket, and the mechanism of action, should also be similar. The results of mutational analysis and functional assays (Heisler et al., 1993; Severinov et al., 1995; Temiakov et al., 2005; Tuske et al., 2005; Yang and Price, 1995) and the structural alignment of *Tth* and *E. coli* ECs (**Suppl Figure S2**) generally support this assumption. The unique structural features of Stl and its mode of interaction with RNAP make it an attractive model for the rational drug design of a new generation of transcription inhibitors (Jeong et al., 2014, 2013).

Despite our knowledge of Stl interactions with RNAP and its inhibitory activities, the exact molecular mechanism of Stl action are not fully understood and remain controversial. The current models inferred from structural analysis and biochemical ensemble assays do not address several important questions. <u>First</u>, what is the role of TL in Stl action? While TL makes several critical contacts with the acetamide group of the Stl tetramic acid moiety (Tuske et al., 2005; Vassylyev et al., 2007) and is essential for the inhibitory function, it appears dispensable for Stl binding to RNAP (Temiakov et al., 2005). <u>Second</u>, does Stl recognize a specific state of RNAP (e.g., an open TL/BH or another conformation it assumes during transcriptional pausing), or is its binding completely stochastic and independent of TL/BH conformation? The former would be consistent with the tight arrangement of the Stl pocket, where the DNA and four mobile RNAP elements must jointly fold to accommodate the inhibitor (Temiakov et al., 2005; Tuske et al., 2005; Vassylyev et al., 2007). However, the conformation-dependent Stl binding would likely result in a sequence-specific pattern of transcript inhibition that has not yet been observed. The alternative mode of Stl action implies that inhibition occurs randomly during elongation, with a certain probability at each NAC. This mode would be consistent with the decrease in the maximum rate of catalysis in single-nucleotide addition reactions observed in ensemble assays (Temiakov et al., 2005; Tuske et al., 2005). <u>Another intriguing question</u> is how does Stl enter the EC? The target binding site in RNAP is not accessible through either primary, secondary, or the RNA exit channels of RNAP in EC due to steric occlusion, respectively, by the downstream DNA duplex and RNA/DNA hybrid, TL, and RNA product (Vassylyev et al., 2007). <u>A related question</u> concerns the observed Mg^2+^-dependence of Stl inhibitory action (Zorov et al., 2014). Is it a unique feature of *Thermus* RNAPs, or do the enzymes from other bacterial species, such as *E. coli*, possess the same trait? <u>Lastly</u>, the interplay between Stl and transcript cleavage factors GreA/GreB is puzzling. Unlike Stl, Gre factors bind RNAP through the secondary channel and only when TL is in the open conformation (Abdelkareem et al., 2019; Roghanian et al., 2011). Also, while TL closing is essential for RNA synthesis, it remains open during the factor-catalyzed endonucleolytic cleavage of the backtracked RNA (Abdelkareem et al., 2019; Sekine et al., 2015). Therefore, GreA/GreB and Stl should not compete for binding to RNAP, and the cleavage reaction should be resistant to Stl. Then what is the molecular basis for the observed inhibition of GreA-induced RNA cleavage in backtracked ECs by Stl (Temiakov et al., 2005)? Does Stl act as an allosteric competitor of GreA, or does it prevent RNA backtracking without affecting the factor binding?

To help address these questions and fill the gaps in our understanding of Stl mechanism of action, we investigated the dynamics of Stl inhibition of *E. coli* RNAP during transcription elongation *in vitro* at a single-molecule level using acoustic force spectroscopy (Sitters et al., 2015). The single-molecule (SM) approaches, including optical trapping (OT), magnetic tweezers, atomic force spectroscopy, and acoustic force spectroscopy (AFS), provide direct real-time measurements of complex multistep stochastic processes, such as transcription, with high spatial (nanometer distance range) and temporal (millisecond time scale) resolution (Abbondanzieri et al., 2005; Dulin et al., 2013; Mohapatra et al., 2020; Revyakin et al., 2006; Sabantsev et al., 2022). Unlike conventional biochemical and biophysical methods, which measure the population-average properties of transcribing RNAP molecules, SM methods assess the properties of individual members of molecular populations, reporting on the functional dynamics of RNAP molecules under defined experimental conditions (Davenport et al., 2000; Herbert et al., 2010; Mejia et al., 2015; Shaevitz et al., 2003; Zhou et al., 2011). Both optical trapping and AFS have been successfully used by us and others to monitor the dynamics of transcription elongation and establish kinetic parameters of the process and characterize the inhibitory activity of three related lasso peptide antibiotics, microcin J25 (Adelman et al., 2004), klebsidin and acinetodin (Metelev et al., 2017). The AFS data revealed that these peptides increase the number and duration of short-lived RNAP pausing events without affecting the instantaneous velocities between the paused states, consistent with the view that the three lasso peptides act by blocking the secondary channel and thus preventing the NTP substrates from accessing the RNAP active site (Metelev et al., 2017).

Here we employed AFS to quantitatively characterize the efficiency of Stl inhibition of RNAP elongation under conditions that promote diverse types of pausing events. We show that Stl does not target the backtracked ECs. Instead, it acts distributively, targeting short-lived (<20 sec) paused ECs that form during elemental pausing, inducing their conversion into long-lived (>20 sec) pausing events. We also demonstrate that the Stl inhibitory action on *Eco* ECs is Mg^2+^-dependent, consistent with previous observations made with *Thermus aquaticus* (*Taq*) RNAP (Zorov et al., 2014). Further characterization of the combined action of Stl and transcript cleavage factors GreA and GreB reveals that Stl does not compete with Gre factors for binding ECs. Instead, GreA and GreB act cooperatively with Stl, amplifying its inhibitory activity during transcription elongation. We propose a plausible structural model of the *Eco* EC-GreB-Stl complex that explains the observed Stl activities and provides insight into possible cooperative action of secondary channel factors and other antibiotics binding at the Stl-pocket.

## MATERIALS AND METHODS

### RNAP and GreA/GreB purification

Biotinylated RNAP was purified from *E. coli* BL21DE3 cells transformed with pIA497 plasmid expressing RNAP β’ subunit fused with biotin carboxyl carrier protein, as described previously (Metelev et al., 2017; Sabantsev et al., 2022). Recombinant GreA and GreB proteins carrying C-terminal 6xHis tag were expressed in *E. coli* Rosetta cells (NEB) transformed with pJL-GreA-CPH and pJL-GreB-CPH plasmids, respectively, and purified by Ni-chelating HisTrap HP (Cytivia) and gel-filtration Superdex 75 10/30 (Cytivia) column chromatography as described (Borukhov and Goldfarb, 1996).

### Stalled complex tether construction

EC carrying 20 nt-long RNA (EC-20) was prepared as described previously (Metelev et al., 2017; Sabantsev et al., 2022) using biotinylated RNAP and a 4413 bp-long DNA template containing T7A1 promoter upstream of *E. coli rpoB* gene followed by two *rrnB* T1 and *rrnB* T2 terminators. Both DNA strands at the downstream end of the template carried digoxigenin-modified nucleotides to allow the immobilization of stalled EC-20 on the AFS chip surface covered with anti-digoxin antibodies (Metelev et al., 2017; Sabantsev et al., 2022). Briefly, 100 nM of biotinylated RNAP and 2 nM DNA template were incubated in transcription buffer (40 mM Tris-HCl, 40 mM KCl, 1 mM DTT, and 5 mM MgCl_2_) for 20 minutes at 37 °C in the presence of 500μM dinucleotide primer ApU (IBA), 35 μM each of ATP, GTP, and CTP. Immediately after EC-20 preparation, it was supplemented with heparin 50 μg/ml and incubated on ice for 1 min.

### AFS experimental setup

Single-molecule (SM) AFS experiments were carried out according to the previously published protocol (Metelev et al., 2017; Sabantsev et al., 2022), the force inside the AFS chip was applied from the bottom side (Kamsma et al., 2016; Kamsma and Wuite, 2018). Commercial AFS chip (Lumicks) contained a special microfluidic chamber with an integrated piezo element on the upper side of the flow cell. Applying voltage to the piezo element produced standing planar acoustic waves generating force applied to the polystyrene microspheres inside the chamber in the direction towards the top side of the chamber The AFS chip was first passivated by treatment with 0.05 mg/ml of anti-digoxigenin antibodies (Roche), followed by incubation with 0.2% BSA (Amresco) and 0.5% pluronic (Pluronic® F-127, Sigma P2443-250G) in PBS for 30 min at 25°C. Next, the biotinylated EC-20 was added to the chamber and immobilized on the surface of the AFS chip via digoxin-labeled DNA. The AFS chip was flushed with a final transcription buffer (40 mM Tris-HCl, 80 mM KCl, 0.5 mM DTT, 10 mM MgCl_2_, 0.02% pluronic, 0.02% casein (Sigma), 50 μg/ml heparin, 1% DMSO (Helicon) followed by addition of 10 μl streptavidin-coated microspheres (2.1 μm) (Spherotech, Cat № SVP-20-5). After washing away the unbound microspheres, the resulting EC-20 immobilized on microspheres via biotinylated RNAP and tethered to the AFS chip via digoxin-labeled DNA was used in transcription elongation experiments as described previously (Metelev et al., 2017; Sabantsev et al., 2022).

4-6 pN force was applied to the microspheres carrying immobilized EC-20 opposite the direction of transcription, and tether formations were observed. The value of the applied force was chosen to minimize its impact on the observed effects (Adelman et al., 2002; Neuman et al., 2003; Shaevitz et al., 2003; Wang et al., 1998) and was kept at below the stalling force (23 pN) or the force that could have a significant adverse effect on transcription (>15 pN) (Adelman et al., 2002; Larson et al., 2011; Neuman et al., 2003; Wang et al., 1998; Zhou et al., 2011). To generate the elongation profiles, we initiated transcription reactions by flowing in transcription buffer containing 1 mM NTPs to the immobilized EC-20 following the movement of RNAP by monitoring the displacement of microsphere bead coordinates using AFS Labview-based software (Lumicks) (Kamsma et al., 2016; Kamsma and Wuite, 2018; Sitters et al., 2015). The experimental scheme is presented in **Suppl. Figure S1**.

In SM-AFS experiments, the following parameters were measured: (i) average velocity of transcription; (ii) instantaneous velocity, i.e., velocity between pauses (>2.5 s); (iii) pause duration (RNAP dwell time); and (iv) average number of pauses (the number of pauses per 500 nm of elongation profile). The elongation profiles of RNAP obtained in the presence of 1 mM of NTPs alone were used as controls.

### SM data analysis

Elongation profiles of individual RNAP complexes were recorded and further analyzed using a custom-made LabVIEW-based program. Only processive RNAP molecules that translocated at least 500 nm along the DNA were selected. For each trace, cut to 500 nm, we calculated the number of pausing events, the duration of each pause, and instantaneous velocity – the rate of pause-free RNAP translocation. The raw data were smoothed by a 15-second median filter followed by Savitsky-Golay filtering (SG rank = 3 s, polynomial order = 3). The instantaneous velocity at each time point was calculated as a derivative. Long pauses were detected as parts of the elongation profile in which the instantaneous velocity did not exceed the preset threshold. To analyze the effect of short pauses, raw data were first filtered to remove the long pauses, then smoothed by a softer Savitsky-Golay filter (SG rank = 0.5 s, polynomial order = 3, LPF rank = 1.5 s), and then used to calculate the instantaneous velocities. Short pauses were detected as instances in which the instantaneous velocity did not exceed the threshold (0.5 s.d.). Pause-free RNAP translocation was further analyzed by plotting the instantaneous velocity as a histogram which revealed two peaks, one centered around zero and another corresponding to the true instantaneous velocity of RNAP. To remove the zero peak, we reflected the distribution profile symmetrically relative to zero and then subtracted it from the original distribution profile.(Arseniev et al., 2022; Bintu et al., 2012; Hodges et al., 2009). The resulting single peak was fitted using Gaussian to calculate the average instantaneous velocity. For each experimental condition, data from multiple individual traces were analyzed, averaged, and presented as mean ± s.e.m. The statistical test to analyze differences in RNAP elongation rates, pause probabilities and average transcription velocities was performed using Mann-Whitney two-tailed tests (significance level p: * ≤ 0.001, ** = ≤ 0.005; *** = ≤ 0.01; **** = ≤ 0.05; n.s. = non-significant).

### Visualization and protein structure analysis

The 3D structures of *Tth* and *Eco* ECs and their complexes with Stl and GreB were superimposed, analyzed, and visualized by ICM-Pro v. 3.9-1b software (Molsoft).

## RESULTS

### Stl induces long-lived pauses

To study the inhibition of transcription elongation of *E. coli* RNAP by Stl at a single-molecule level, we utilized the AFS technique (Metelev et al., 2017; Sabantsev et al., 2022; Sitters et al., 2015). When the initial EC-20 was extended under standard reaction conditions in the presence of 1 mM NTPs (saturating concentrations) and 10 mM MgCl_2_, the elongation profiles exhibited a monotonous growth of transcript length over time with a few short-lived (<10 sec-long) pauses and no long-lived pauses (**Figure 1A, 1B**).

**Figure 1.**
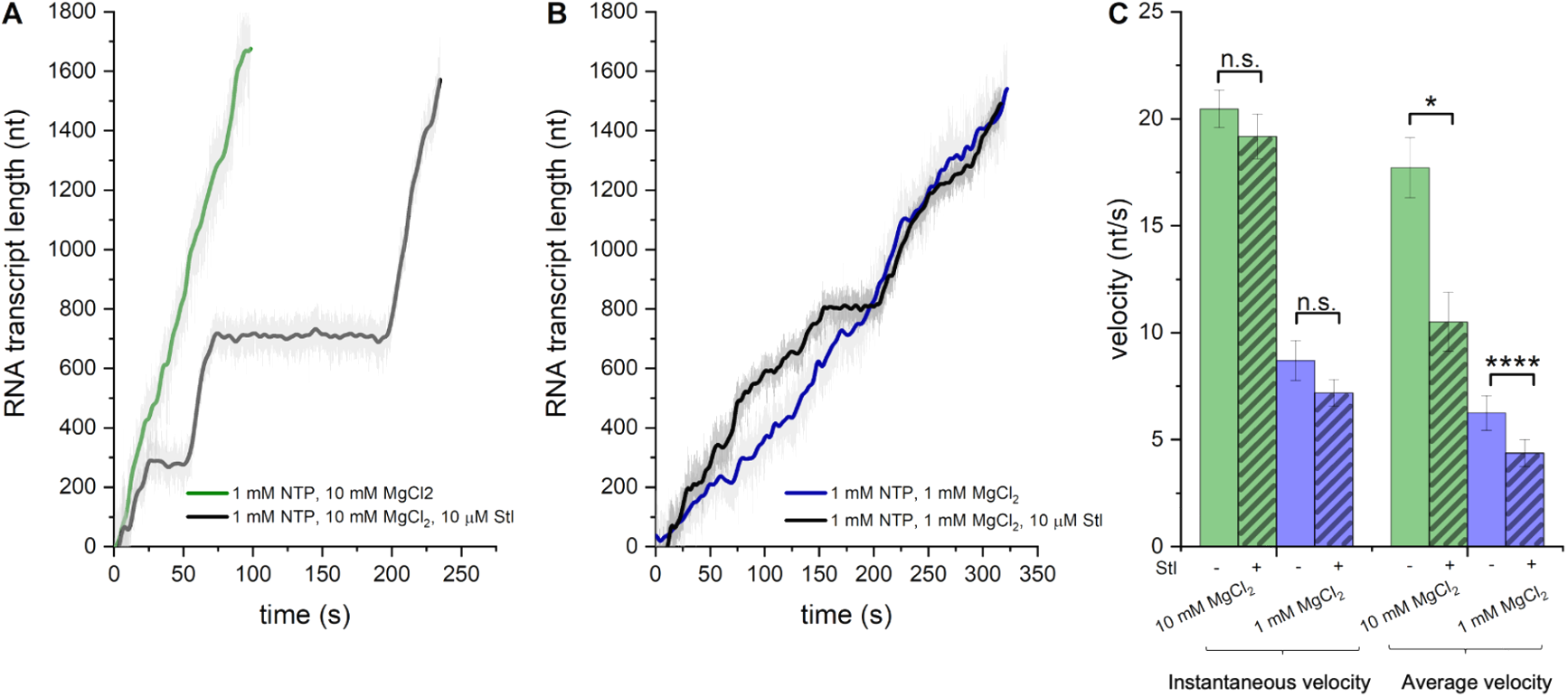
Stl inhibits transcription elongation by inducing long-lived pauses in a Mg^2+^-dependent manner. **A, B**. A single-molecule AFS experiment showing representative elongation profiles (plotted as RNA nucleotides transcribed vs. time) recorded for individual RNAP molecules at 1 mM NTPs in the absence (green and blue traces) and presence (gray and black traces) of 10 μM Stl at 10 mM Mg^2+^ (**A**, 28 and 33 traces, respectively) and 1 mM Mg^2+^ (**B**, 9 and 11 traces, respectively). **C**. Bar graph showing the instantaneous (green and blue bars) and average (brown and red bars) velocities for individual RNAPs during transcription in the absence (non-hatched bars) and presence (hatched bars) of 10 μM Stl at 10 mM Mg^2+^ (left panel) and 1 mM Mg^2+^ (right panel). Data were calculated based on the analysis of multiple elongation profiles P-value for all cases are indicated as: * = ≤ 0.001; ** = ≤ 0.005; *** = ≤ 0.01; **** = ≤ 0.05; n.s. = non-significant.

The observed kinetic parameters, including instantaneous and average velocities, were in good agreement with the results reported previously under similar conditions (Adelman et al., 2004; Janissen et al., 2022; Mejia et al., 2015; Metelev et al., 2017; Shaevitz et al., 2003). The addition of 10 μM Stl to the reaction resulted in the appearance of random long-lived (>20 sec) pausing events (**Figure 1A**). Although rare (1±0.2 occurrences per trace), these pauses noticeably decreased the average velocity of elongation (**Figure 1B**). Yet, the slope of elongation profiles between the pauses was similar to that observed in the control reaction (**Figure 1A**). As a result, the negative effect of Stl on instantaneous velocity was negligible (∼1.1-fold) (**Figure 1B**). Analysis of multiple elongation profiles recorded under these conditions revealed that the average duration of Stl-induced pause was 192 ± 44 seconds (mean ± s.e.m.). Note that the Stl concentration in these experiments was significantly lower than the saturating concentrations (1-3 mM) used for structural studies (Temiakov et al., 2005; Tuske et al., 2005; Vassylyev et al., 2007). The concentration used in this study is slightly below the Stl binding and inhibitory constants towards the *E. coli* RNAP (*K*_*D*_≈15 μM (Tuske et al., 2005); *K*_*i*_ (IC_50_) ≈24 μM (Temiakov et al., 2005)) and thus enables the detection of single acts of inhibition.

### Stl inhibition is Mg^2+^-dependent

We examined whether the inhibitory activity of Stl towards *E. coli* RNAP depends on the concentration of Mg^2+^, similar to that reported previously for *Taq* RNAP (Zorov et al., 2014). To this end, we compared the elongation profiles of *E. coli* RNAP obtained at 10 mM and 1 mM MgCl_2_ in the absence and presence of 10 μM Stl (**Figure 1A, B**). As expected, at a low concentration of Mg^2+^, the instantaneous and average transcription velocities were almost equal to each other and ∼2.5-times lower than under standard reaction conditions (**Figure 1A, B**). However, the inhibitory effect of Stl on the total number and duration of longlived pauses and the average elongation velocity was more substantial at a high concentration of Mg^2+^ (**Figure 2B**). Thus, for both *Thermus* and *E. coli* RNAPs, Stl-inhibition is Mg^2+^-dependent.

**Figure 2.**
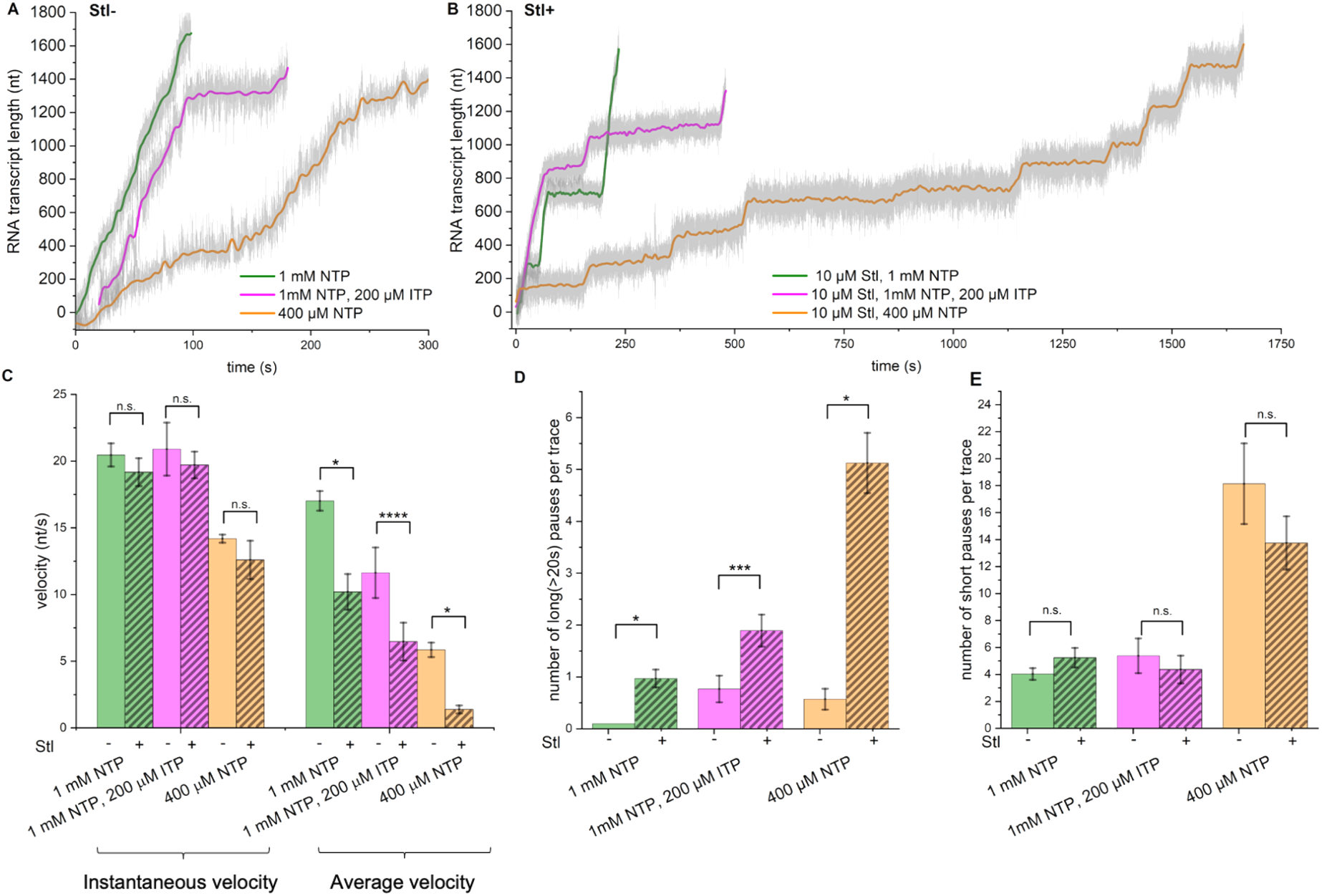
Inhibitory effect of Stl on RNAP pausing intermediates. **A, B**. Representative elongation profiles for individual RNAPs observed with 1 mM NTPs (green), 400 μM NTPs (orange), or 200 μM ITP and 1 mM NTPs (magenta) in the absence (**A**) (N=28, N=7, N=13) or in the presence (**B**) (N=33, N=8, N=19) of 10 μM Stl. Bar graphs showing the instantaneous and average velocities (**C**), the mean number of long (>20 sec) pauses (**D**), and short (2,5–20 sec) pauses (**E**) per elongation profile, calculated from the analysis of multiple elongation traces observed under conditions shown in **A** and **B**. P-value for all cases are indicated as *≤0.001; **≤0.005; ***≤0.01; ****≤0.05; n.s. = non-significant.

### Stl targets RNAP in an elementary pause state

To bind RNAP and inhibit its catalytic activity, Stl may recognize a specific RNAP conformation during transcription, such as an open TL/BH state that occurs every NAC and induces elemental pausing (Kang et al., 2019) or a ratcheted/swiveled state that occurs when RNAP backtracks and long-lived pauses are formed (Abdelkareem et al., 2019; Tagami et al., 2010). Alternatively, Stl binding could be stochastic and independent of RNAP conformation. To explore these possibilities, we correlated the efficiency of Stl inhibition with two types of transcriptional pausing (and the RNAP conformational states associated with these pauses) by manipulating the compositions of NTPs. First, we conducted AFS experiments in the presence of inosine triphosphate (ITP), a non-canonical purine nucleotide substrate that forms weak base pairs with both thymine and cytosine bases in DNA. Upon incorporation into the nascent transcript, inosine destabilizes the RNA/DNA hybrid in ECs, inducing RNAP backtracking and long-lived pausing (Mejia et al., 2015; Shaevitz et al., 2003).

Transcription in the presence of a mixture of 1 mM NTPs with 200 μM ITP resulted in mostly single random long-lived pauses (due to increased backtracking) with an average duration of 123±45 sec (**Figure 2A**), causing a ∼1.5-fold decrease of the average elongation velocity relative to the control. At the same time, the instantaneous velocity and the number of short-lived elemental pauses were unaffected (**Figure 2C-E**). These results are in good agreement with the published data (Mejia et al., 2015; Shaevitz et al., 2003). Reaction in the presence of 10 μM Stl, 1 mM NTPs, and 200 μM ITP led to additional long-lived pauses (**Figure 2D**), causing a further decrease in the average velocity relative to the reactions with either Stl alone or ITP+NTPs (**Figure 2A**). The instantaneous velocity and the number of short-lived (<20 sec) pauses remained unchanged (**Figure 2C-E**). Notably, the number of observed long pauses was roughly equal to the sum of pauses induced by Stl and ITP individually. Thus, Stl does not prolong the backtracked pauses induced by ITP, indicating that Stl and ITP act independently, and their effect on the number of long-lived pauses is additive. From this, we conclude that Stl does not target backtracked paused ECs.

We next investigated whether Stl acts by targeting active ECs that cycle between pre-translocated and post-translocated states or targets transiently paused translocation intermediates, the elemental paused ECs, that are thought to be trapped in a half-translocated state (Kang et al., 2019). To this end, we conducted SM-AFS experiments using reduced concentrations of natural NTPs (Abbondanzieri et al., 2005; Forde et al., 2002; Janissen et al., 2022; Mejia et al., 2015). Transcription under these conditions increases the RNAP dwelling time at each nucleotide position in the DNA template during NAC, promoting TL/BH opening (Mazumder et al., 2020) and stimulating the formation of short-lived elemental pauses (Janissen et al., 2022).

Reducing the concentration of NTPs from 1 mM to 400 μM in the absence of Stl led to a noticeable decrease in both instantaneous and average velocities, apparently due to a sharp (∼4.5-fold) increase in the number of short-lived elemental pauses (**Figure 2A, C, E**). The addition of Stl to the reaction with lowered NTP concentrations resulted in a dramatic increase in the number of long pauses (∼9-fold), causing a ∼3-fold decrease in the average elongation velocity (**Figure 2B-D**). Thus, the reduced concentration of NTPs (below or near the apparent Km) stimulates the inhibitory activity of Stl. We also noted that under these conditions, the number of short pauses was noticeably lower than in the absence of Stl (**Figure 2E**), suggesting that Stl converts a fraction of short pauses into long ones. These results indicate that Stl acts primarily on stalled ECs during elemental pausing or at some intermediate steps en route to a pause state in which RNAP has an open conformation of TL/BH.

### Transcript cleavage factors GreA and GreB enhance Stl inhibitory activity

Our next goal was to investigate the functional interplay between Gre factors and Stl during transcription elongation using the SM-AFS technique. Specifically, we sought to address the long-standing question of whether Stl acts as an allosteric competitor of GreA/GreB. GreA and GreB transiently bind both active and backtracked/paused ECs with similar affinities (K_DGreA_ ∼1 μM and K_DGreB_ ∼100 nM, respectively), occupying the secondary channel of RNAP with their N-terminal coiled-coil domain (Gre-NTD) (Tetone et al., 2017). In paused backtracked ECs, the acidic tip of Gre-NTD, which coordinates Mg^2+^- and OH^-^-ions, reaches the enzyme’s catalytic center and stimulates the cleavage of nascent RNA 3’-tail extruded through the secondary channel (Borukhov et al., 1993; Sosunova et al., 2013). GreA excises 2-3 nt from 3’-end, whereas GreB cleaves out fragments up to 15 nucleotides (Erie et al., 1993; Kulish et al., 2000; Laptenko et al., 2003; Shaevitz et al., 2003). After the cleavage, RNAP resumes transcription from the newly generated RNA 3’-OH that is positioned in the active site. By this “cleavage and restart” mechanism, Gre factors prevent and alleviate backtracked pausing and arrest when RNAP misincorporates or encounters a roadblock to elongation (Borukhov et al., 1993; Borukhov and Goldfarb, 1996; Laptenko et al., 2003). Generally, both factors stimulate transcription *in vitro* and *in vivo*, improving bacterial cell fitness (Erie et al., 1993; Laptenko et al., 2003; Marr and Roberts, 2000). Structural data revealed that in EC-GreA/B complexes, the RNAP TL remains in an open ordered conformation, allowing passage of Gre-NTD through the secondary channel (Abdelkareem et al., 2019; Sekine et al., 2015), temporarily inactivating the enzyme catalysis. Consequently, binding Gre factors to active ECs slows down RNA synthesis (Laptenko et al., 2003; Tetone et al., 2017). However, SM fluorescence microscopy experiments showed that the inhibitory effect of GreB on elongation rates was moderate (<2.5-fold), owing to its highly transient and short-lived association with RNAP even under saturating concentrations (Tetone et al., 2017).

Consistent with these observations, instantaneous and average velocities of EC elongation in the presence of 1 mM NTPs and 5 μM GreA or GreB measured in our SM-AFS experiments were reduced relative to control (20.5 ± 0.87 nt/s) only by ∼10% (18.4 ± 1.4 nt/s) and 55% (9.1 ± 0.7 nt/s), respectively (**Figure 3A, C**). The effects of each factor on average and instantaneous velocities were similar, as neither GreA nor GreB induced long-lived pauses (**Figure 3A, D**). At the same time, the number of short-lived pauses (within the range of 2.5-20 sec detectable by SM-AFS) increased only by 1.5-fold with GreB and was even lower (by 2-fold) with GreA (**Figure 3E**). The observed decreases in instantaneous and average velocities could result from the accumulation of very short (<2.5 sec) pauses induced by GreA and GreB (that are undetectable by AFS) or by changes in the rate of catalysis.

**Figure 3.**
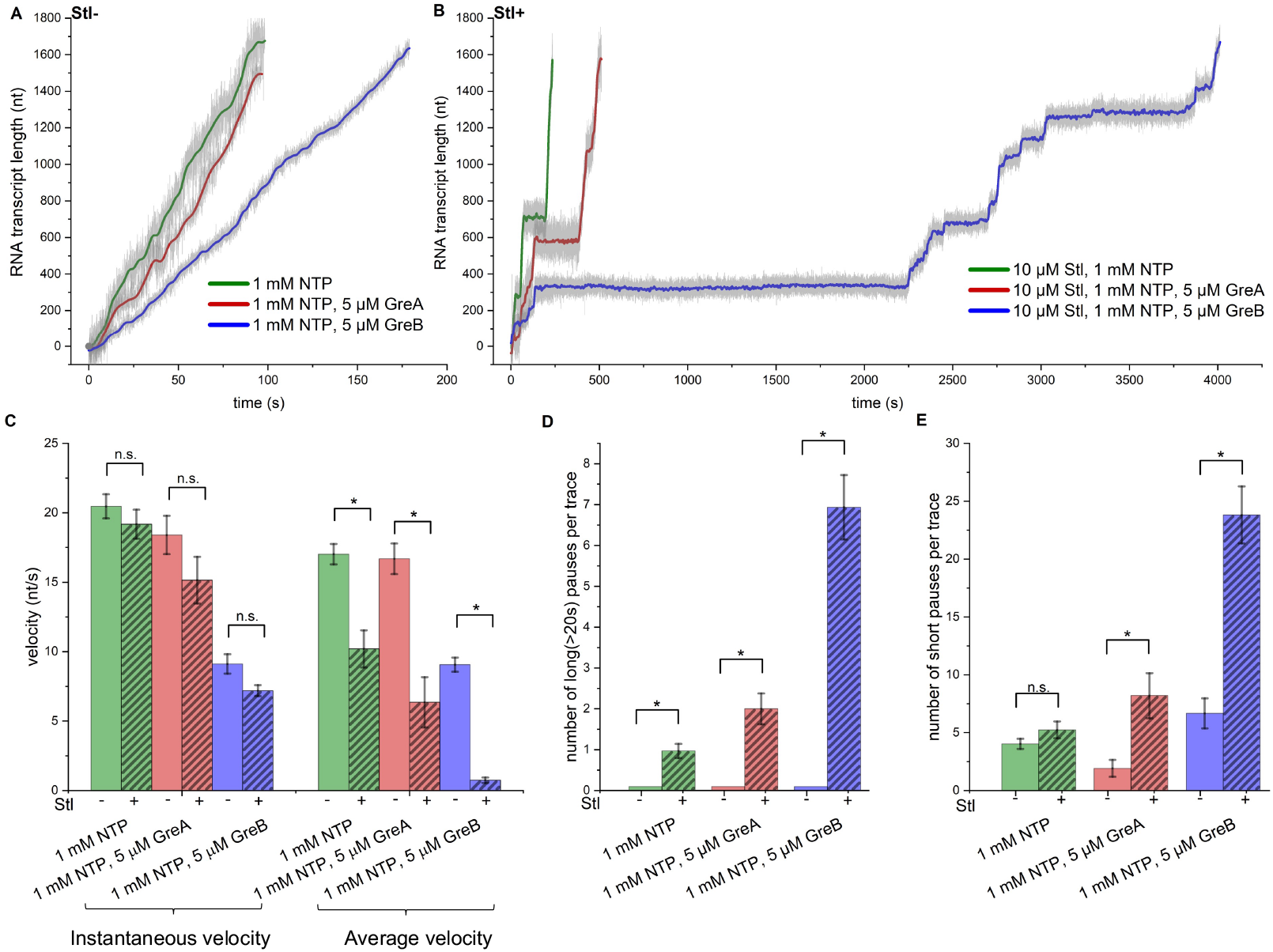
The combined effect of Stl and Gre factors on transcription elongation velocities and pausing. **A, B**. Representative elongation profiles for individual RNAPs observed with 1 mM NTPs (green), 1 mM NTPs and 5 μM GreA (red), 1 mM NTP and 5 μM GreB (blue) in the absence (**A**) N=28, N=11, N=15) and in the presence of 10 μM Stl (**B**) (N=33, N=15, N=16). Bar graphs showing the instantaneous and average velocities (**C**), the number of long-lived (>20 sec) pauses **(D**), and short (2,5–20 sec) pauses **(E**) per elongation profile, calculated from the analysis of multiple elongation traces under conditions shown in A and B. P-value for all cases are indicated as: * = ≤ 0.001; ** = ≤ 0.005; *** = ≤ 0.01; **** = ≤ 0.05; n.s. = non-significant.

Unexpectedly, the combined action of 5 μM GreB and 10 μM Stl in the transcription reaction resulted in a much more significant decrease in the average velocity (∼17-fold relative to the control) than the effects from each component added separately (**Figure 3A-C**). The change in the instantaneous velocity was less extensive (∼2.6-fold) and mainly due to the effect of GreB alone. The drop in the average velocity correlated with the drastic increase (7-fold) in the number of long-lived pauses, their duration (up to 3000 sec), and the number of short-lived pauses (6-fold), relative to the action of Stl alone (**Figure 3 B-E**). Similar results were observed when 5 μM GreA and 10 μM of Stl were added to the transcription reaction (**Figure 3A-E**). The average elongation velocity decreased ∼3-fold relative to the control, owing to the ∼2-fold increase in the number of long-lived pauses (**Figure 3A-D**). In contrast, the instantaneous velocity and the number of short-lived pauses did not change substantially (**Figure 3C, E**). Thus, the combined inhibitory effect of Stl with either GreA or GreB on the average velocity is more synergistic than additive. We also conclude that Stl does not compete with Gre factors for binding to RNAP during transcription elongation. Instead, GreA and GreB act cooperatively with Stl, effectively amplifying its inhibitory activity.

### The structural model of the *Eco* EC-GreB-Stl complex explains the synergy between Gre factors and Stl

To better understand the mechanistic basis for the synergistic effects of GreA and GreB on Stl inhibition, we superimposed the high-resolution structures of *Tth* EC-Stl inhibitory complex (PDB: 2PPB; (Vassylyev et al., 2007) and *Eco* ECs with the bound substrate (PDB: 6RH3) and GreB (PDB: 6RI7) (Abdelkareem et al., 2019). Consistent with the earlier predictions (Temiakov et al., 2005; Tuske et al., 2005), the structural alignments of *Tth* and *Eco* ECs revealed that the spatial arrangement of the key mobile elements (including TL, BH, FL2, and βD-loopII), forming the Stl-binding pocket, and most of the hydrophobic and polar contacts with Stl are conserved between *Tth* and *Eco* enzymes (**Suppl. Figure S2A**). The notable exceptions include the presence of *Eco* β’N792 (instead of *Tth* β’D1090) in the BH element, which reduces the Stl binding affinity to *Eco* RNAP (Temiakov et al., 2005), and *Eco* β’A1142 (instead of *Tth* β’I1260) in TL, the side chain of which is too small to contact the methylacetamide group in the tetramic acid moiety of Stl (**Suppl. Figure S2B**). Otherwise, the superimposed Stl molecule firmly fits into *Eco* RNAP antibiotic-binding pocket without any clashes with the side chains of its residues.

Compared to the structure of *Eco* EC in the translocated state with the bound substrate, GreB insertion into RNAP secondary channel induces rearrangements of several domains (Abdelkareem et al., 2019). These include (i) an upward swiveling movement of the β’ rim helices domain (by ∼7Å) together with the F-loop, resulting in channel expansion, (ii) an inward movement of β lobe2 with βSI1 domain towards the active center (by ∼5Å), pushing three β subunit loops FL2 (residues T530-F545), D-loopII (P564-G570), and β lobe2 loop (D160-L171) closer to the Stl-binding pocket (by ∼2Å), and (iii) most importantly, an outward swiveling movement of the TL (that adopts an open conformation) together with the β’SI3 domain, resulting in the further opening of the secondary channel and additional tightening of the Stl-binding pocket (**Suppl. Figure S3**). Superimposing the two *Eco* EC structures with the *Tth* EC-Stl inhibitory complex structure shows that GreB-induced domain rearrangements in RNAP leave the Stl-binding pocket largely unaltered (within a margin of 1-1.5Å). A notable exception is the induced movement of TL and βD-loopII that brings five more residues into proximity to the Stl tetramic acid moiety, possibly strengthening its binding to RNAP. These include TL β’M932 contacting acetamide group, and TL residues β’R933, β’F935, β’I1134, and β lobe2 loop residue βK163 interacting with 3-hydroxy-2-methyltetrahydropyranyl group **(Suppl. Figure S2B, S3, and Figure 4A**). Furthermore, a comparison of the *Eco* EC-Stl and *Eco* EC-GreB-Stl model structures indicates that the only entry route for Stl into RNAP is the narrow channel formed by TL, F-loop, FL2, βDloopII, jaw domain, the rim helices tip, and the downstream dsDNA held in place by the β’ clamp (**Suppl. Figure S3)**. In the model, GreB does not interact with Stl directly; instead, it contributes to Stl binding allosterically by further constricting the Stl-entry channel (primarily via repositioning of open TL), thereby preventing Stl dissociation from EC (**Figure 4B)**. As a result, Stl becomes trapped inside RNAP, thus dramatically increasing its inhibitory potency.

**Figure 4.**
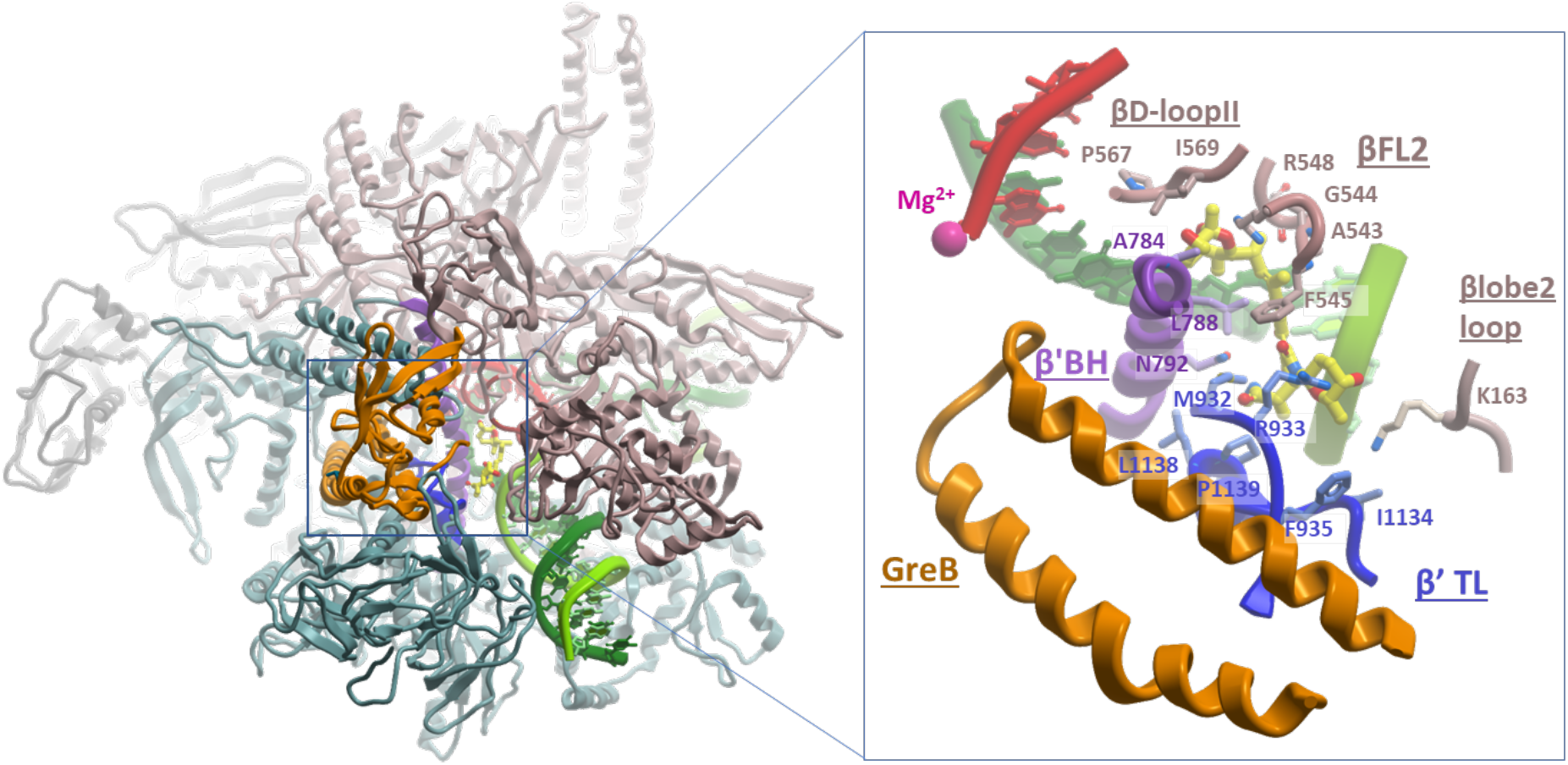
Proposed structural model explaining the synergistic effect of GreB on Stl binding in *Eco* EC. **(A)** Structural model of *Eco* EC-GreB-Stl complex shown as colored ribbons: αI and αII, light gray/gray; β, coral; β’, teal; GreB, orange. The active center mobile elements, TL, BH, FL2, β D-loopII, and β lobe2 loop are colored blue, purple, coral, and brown, respectively. The nucleic acids are shown as colored ribbons and sticks: RNA, red; DNA template strand, dark green; and DNA nontemplate strand, green. Stl is shown as CPK-colored sticks (except carbon atoms are in yellow), and the catalytic Mg^2+^ is shown as a magenta ball. The inset on the right (blue rectangle) shows the zoom-in view of the selected area near the Stl-binding pocket. The inset view is obtained by rotating the left view by ∼40° around the vertical axis. Residues of RNAP mobile elements contacting Stl are indicated. **(B)** Comparison of the Cryo-EM structures of *Eco* post-translocated EC with bound CTP substrate (PDB: 6RH3, left) and *Eco* EC-GreB (PDB: 6RI7, right) shown as solvent-accessible surfaces with superimposed Stl (CPK-colored balls). The surfaces of RNAP subunits are colored as in (A). The insets below show the magnified view of the Stl-entry channel in each structure.

## DISCUSSION

In this work, we applied the SM-AFS technique to investigate the targeting and the dynamics of Stl inhibition of *E. coli* RNAP transcription elongation using a well-characterized 4413 bp-long DNA template containing the *rpoB* gene (Dangkulwanich et al., 2014; Larson et al., 2011; Metelev et al., 2017; Sabantsev et al., 2022; Schafer et al., 1991; Shaevitz et al., 2003). We made several important observations that offer new insights into the mechanism of Stl inhibition and its synergism with transcription factors GreA and GreB. First, we showed that under subsaturating concentrations (10 μM), Stl induces rare long-lived pausing events that appear randomly distributed along the DNA (**Figure 1A**). At the same time, Stl does not affect the instantaneous velocity of elongation (**Figure 1C**), indicating that it does not act in the on-pathway of NAC. We also found that Stl action requires a high concentration (10 mM) of Mg^2+^ ions (**Figure 1**), similar to an earlier observation with *Taq* RNAP (Zorov et al., 2014). Thus, dependence on noncatalytic Mg^2+^ for Stl inhibition is a common feature of RNAPs from two diverse bacterial species. However, *Eco* RNAP carries BH residue β’N795 at the same position as β’D1090 in *Taq* RNAP and, therefore, cannot coordinate Mg^2+^ ion, as hypothesized by Zorov et al., 2014. Therefore, we propose an alternative explanation for the observed Mg^2+^ dependence of Stl inhibition. The noncovalent Mg^2+^ ion may bridge the Stl tetramic acid carbonyl/hydroxyl groups and nontemplate DNA strand i+3/i+4 phosphate, thus stabilizing the Stl-EC inhibitory complex (see **Suppl Figure S2B**).

Next, we showed that Stl does not affect the long-lived backtracked pauses induced by the incorporation of ITP into nascent RNA (**Figure 2**), indicating that Stl does not target ECs in the off-pathway backtracked state. These results suggest that Stl binds inefficiently to the active post-translocated state ECs or the paused backtracked ECs. Instead, we found that transcription under reduced NTP concentration, which promotes the formation of short-lived pauses, stimulates Stl-induced long-lived pausing (**Figure 2**), thereby exacerbating its inhibitory action. Notably, the accumulation of the long-lived pauses is accompanied by a proportional reduction of the short-lived pausing events (**Figure 2**). These observations lead us to the conclusion that Stl preferentially targets the off-pathway short-lived paused ECs, where RNAP exists in a catalytically inactive elemental pause state (Landick, 2006).

Lastly, our investigation of the functional interplay between Stl and transcript cleavage factors GreA and GreB led to an unexpected result. We found that instead of displaying competitive behavior, both Gre factors acted cooperatively with Stl, dramatically stimulating the number and duration of long-lived pausing events, albeit to a various degree (**Figure 3**). Remarkably, the combined inhibitory effect of Gre/Stl on transcription elongation was more synergistic than additive, especially for GreB (**Figure 3C-E**). These observations are unusual and somewhat puzzling, considering that Stl strongly inhibits the Gre-induced nucleolytic reactions in backtracked ECs (Temiakov et al., 2005). To our knowledge, this is the first example of synergism between an antibiotic and a transcription factor.

The results of our SM-AFS experiments appear to be in disagreement with the previous biochemical data demonstrating that Stl can inhibit all enzymatic activities of *Tth* and *Eco* RNAPs, including single-nucleotide addition reactions, pyrophosphorolysis, and nucleolytic hydrolysis of nascent RNA in backtracked ECs by reducing the maximal rate of catalysis (Temiakov et al., 2005; Zorov et al., 2014). These observations imply that Stl can bind active pre-and post-translocated ECs and paused backtracked ECs.The X-ray crystal structure of the *Tth* EC in a post-translocated state in complex with Stl and the substrate analog also supported this view (Vassylyev et al., 2007). The apparent discrepancy can be explained by the difference in the experimental setup. In previous studies, stalled ECs formed on RNA/DNA scaffolds were pretreated with Stl used at saturating concentrations (166 μM-1 mM) prior to incubation with substrates in the single-nucleotide extension, pyrophosphorolytic, and transcript cleavage reactions (Temiakov et al., 2005; Zorov et al., 2014). Cryo-EM and SM-FRET data accumulated to date indicate that ECs exist in dynamic equilibrium between multiple conformational states (Kang et al., 2019; Mazumder et al., 2021; Saba et al., 2019) that may have different translocation registers of the RNA/DNA hybrid (such as the half-translocated state seen in elemental paused ECs, ePECs), folded/unfolded (closed/open) conformations of TL, rotational/swiveling states of β’ rim helices, β’SI3 and β’ clamp domains (Abdelkareem et al., 2019; Kang et al., 2019; Murakami, 2015; Saba et al., 2019; Sekine et al., 2015). Some of these conformations could be more susceptible to Stl than others (see below). Preincubation of stalled ECs with Stl (Temiakov et al., 2005; Zorov et al., 2014) substantially increases the time window for Stl binding not only to ECs in the preferred conformational state (which initially may represent only a small fraction of all states) but also to other, less favorable, states. Indeed, ECs treatment with Stl prior to the addition of substrates increased the efficiency of Stl inhibition of different enzymatic activities of RNAP (Zorov et al., 2014). In contrast, our SM-AFS experiments use a continuous transcription elongation system where NTPs are added to the reaction together with Stl at sub-saturating (10 μM) concentrations. These Stl concentrations are comparable to its MIC of ∼5 μM for susceptible *E. coli* strains and are more physiologically relevant (Deboer et al., 1955; Tuske et al., 2005). Under these conditions, the time window for Stl binding would be very small, probably in the millisecond range, allowing only the bona fide targets to be recognized and acted on.

The entry of active ECs into the elemental pause state during NAC is triggered by sequence-specific interactions of RNAP with DNA template known as universal consensus pausing sequence (CPS) and can be further modulated by the presence of nascent RNA secondary structures (e.g., pause-inducing RNA hairpins), cis- and trans-acting protein factors (NusA, NusG, RfaH, Nun, etc.), and small-molecule regulators (ppGpp) (Kang et al., 2019). The spatial resolution of the AFS technique is insufficient to identify the exact position of the Stl-induced pauses in the DNA template used in the experiments. However, with the average frequency of CPSs of 1 per ∼100 bp (Herbert et al., 2006; Landick, 2006; Neuman et al., 2003; Vvedenskaya et al., 2014), we anticipate ∼15-20 potential elemental pausing sites on our template. Considering the apparent Kd ≈15 μM for the Stl-RNAP complex (Tuske et al., 2005) and that RNAP may not pause at each CPS site, the estimated number of short-living elemental pauses induced by 10 μM Stl (in case its binding to ECs is not selective) should be ∼3-5/trace, which is significantly higher than the observed 1-2 pauses/trace. At the same time, we do not detect any discernable pausing pattern in the 33 elongation traces analyzed (**Figure 1**). Because elemental pauses have a typical lifetime of several seconds both *in vivo* and *in vitro* (Churchman and Weissman, 2011; Davenport et al., 2000; Herbert et al., 2006; Larson et al., 2014, 2011; Neuman et al., 2003), we propose that Stl stochastically binds only to a fraction of elemental paused ECs (ePECs).

What could be the structural basis for the observed Stl preference for ePECs? Examination of the Cryo-EM structure of the active *Eco* EC captured in a post-translocated state with the bound substrate, closed TL, and rotated β’SI3 domain (PDB: 6RH3, (Abdelkareem et al., 2019) reveals that the Stl-entry channel is too narrow (Ø≈16-18Å) to allow free diffusion of Stl molecule which has a comparable size (8×18 Å) (**Suppl. Figure S4**). Therefore, Stl binding would be impeded, explaining why the active post-translocated EC in such a closed conformation would be a poor target for Stl. However, a recent Cryo-EM study of *Eco* ePECs formed on CPS templates(Kang et al., 2023) revealed that a significant fraction (45%) of the observed conformational intermediates exists in a pre-translocated state with open unfolded TL and outward rotated β’SI3 domain and β’ rim helices (*con*-ePEC_ufTL, PDB:8EGB). In this conformation, the Stl-entry channel becomes much wider (Ø≈25-40Å) than observed in the active post-translocated state *Eco* EC, allowing free passage of the Stl molecule to its binding pocket (**Suppl. Figure S4**). Therefore, we propose that such intermediate state ePECs serve as specific primary targets for Stl. In subsequent steps, Stl would dock to its binding pocket deep inside the channel, and interact with TL, inducing its partial folding and stabilizing its open-state conformation. This would prevent TL from reaching the catalytic site, effectively inhibiting all RNAP enzymatic activities. Curiously, the Stl-entry channel in the structures of *Eco* backtracked EC both in swiveled (PDB: 6RIP) and not-swiveled (PDB: 6RI9) conformations (Abdelkareem et al., 2019) is also wide open enough to accommodate Stl binding with subsequent stabilization of open state TL. This is consistent with the observed Stl inhibition of the intrinsic nucleolytic activity of RNAP in backtracked ECs (Temiakov et al., 2005; Zorov et al., 2014). We hypothesize, however, that as the Stl-entry channel remains open in backtracked ECs, Stl dissociates before it can extend the paused state, which explains its negligible effect on ITP-induced long-lived pauses (**Figure 2**).

Our structural model of the EC-GreB-Stl complex (**Figure 4**) shows how GreB (and, by analogy, GreA) locks Stl inside RNAP, preventing it from dissociation and providing a reasonable mechanistic basis for the observed synergism between Gre factors and Stl during pausing. The synergistic effect of GreB on Stl inhibititor activity is much stronger than that of GreA, apparently due to its higher binding affinity to RNAP (Laptenko et al., 2003). The model implies that Stl must enter the ePEC first, followed by the binding of GreA/B. The interactions of the Gre factor with ECs are fast and highly dynamic (Tetone et al., 2017), making this scenario plausible. However, if Gre and Stl do not compete and can jointly bind ECs, how can Stl exert its inhibitory effect on GreA/GreB-activated RNA cleavage? Although TL in the EC-GreB complex assumes an open conformation and is not involved directly in catalysis, it contributes to the proper placement of the N-terminal coiled-coil domain of Gre protein (Gre-NTD) in the secondary channel (Abdelkareem et al., 2019; Laptenko et al., 2003), and is essential for the efficient cleavage reaction (Miropolskaya et al., 2017). Because Stl binding alters the TL conformation, we hypothesize that this would compromise the positioning of Gre-NTD catalytic residues at the active center of RNAP, thus interfering with Gre-induced RNA hydrolysis.

Our discovery of cooperativity between Gre factors and Stl in transcription inhibition has several potential implications for the understanding of transcription regulation and for biomedical applications. First, other transcription factors (e.g., DksA, TraR, and Rnk) and inhibitors (e.g., lasso peptides microcin J25, klebsidin, and capistruin and Salinamide-A) that act through RNAP secondary channel also interfere with TL folding (directly or indirectly) and thus may alter the lifetime of ePEC and amplify the pause-inducing activity of Stl. On the other hand, the secondary channel factors may potentiate the inhibitory activity of other small-molecule inhibitors binding in the Stl-entry channel and adjacent pockets, including CBR antimicrobials, Salinamide-A, and GE23077 (Bae et al., 2015; Degen et al., 2014; Zhang et al., 2014). Experiments are currently on the way to test these hypotheses *in vitro* and *in vivo*. Finally, these results may also point to a more effective strategy for the high-throughput screening of new antimicrobial agents. Instead of employing RNAP as a lone target in the inhibitory assays, we propose to use an array of RNAP complexes with various transcription elongation factors essential for bacterial physiology, such as GreA/B, DksA, NusA, NusG, Mfd, UvrD, etc. These factors alter the conformational states of RNAP during transcription, thus presenting more relevant targets and increasing the chances of selecting and identifying more potent inhibitors.

## FUNDING

**Ministry of Science and Higher Education of the Russian Federation under the program “Priority 2030”, 075-15-2021-1333, dated 30.09.2021**

Mikhail Panfilov, Georgii Pobegalov, Alina Potyseva, Maria Yakunina, Mikhail Khodorkovskii

**Ministry of Science and Higher Education of the Russian Federation (project no. 075-15-2021-1062)**

Anatolii Arseniev

**The intramural funds from the Department of Cell Biology and Neuroscience at Rowan University**

Sergei Borukhov

**National Institute of Health Grant R01GM13094**

Sergei Borukhov

**Grant from the Ministry of Science and Higher Education of the Russian Federation (agreement No. 075-10-2021-114 from 11 October 2021)**

Konstantin Severinov

## CONFLICT OF INTEREST

None declared

## Notes

### Competing Interest Statement

The authors have declared no competing interest.

