## Supplemental files for "Single-molecule studies reveal the off-pathway elemental pause state as a target of streptolydigin inhibition of RNA polymerase and its dramatic enhancement by Gre factors"

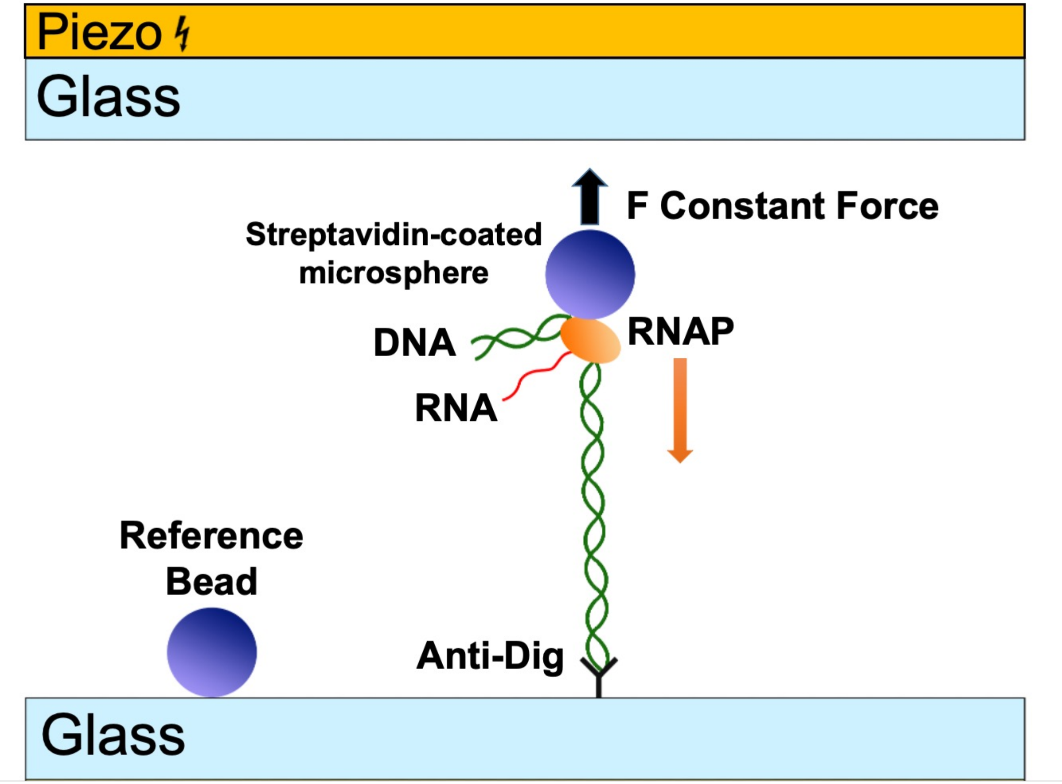


**Suppl. Figure S1.** SM-AFS experimental setup. Elongation profiles of individual transcribing RNAP complexes were obtained using a commercial AFS setup (Lumicks B.V., Amsterdam). The digoxigenin-modified downstream end of the DNA molecule was attached to the anti-digoxigenin-coated surface of the AFS chip. A streptavidin-coated polystyrene bead was attached to a stalled biotinylated RNAP. A tether was stretched with a constant force of 4-6 pN by applying a voltage to the piezoelectric element. Transcription elongation was resumed by supplying the AFS chip with NTPs, and the RNAP translocation dynamics was registered by tracking the position of the bead in 3D.

A

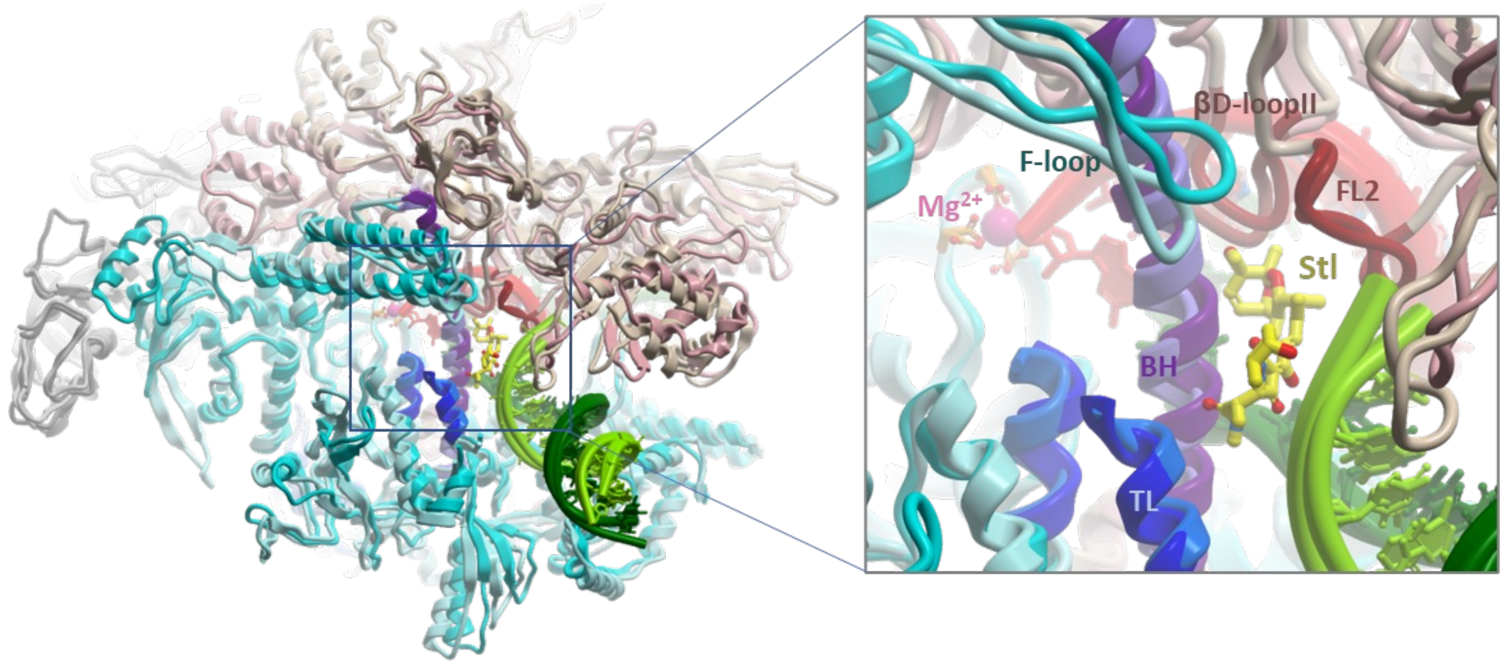


**Suppl. Figure S2. (A).** Structural alignment of Tth EC-Stl (PDB: 2PPB; Vassylyev et al., 2007) and Eco EC (PDB: 6RH3; Abdelkareem et al., 2019) complexes. The view is depicted from the RNAP secondary channel side and the downstream dsDNA-binding cleft. The diagram on the right is a zoom-in active center view of the structures on the left. The structures are shown as colored ribbons: αI and αII, light gray/gray; β, tan/pink; β', aqua/cyan (lighter colors, Tth RNAP; darker colors, Eco RNAP). The active center mobile elements, TL, BH, and FL2, are colored light blue/blue, lavender/purple, and dark red/brown, respectively. The nucleic acids are shown as colored ribbons and sticks: RNA, red; DNA template strand, dark green; and DNA nontemplate strand, green. The Stl and the catalytic center aspartates are shown as CPK-colored sticks (except carbon atoms are in yellow), and the catalytic Mg2+ is shown as a magenta ball. Several domains in both RNAPs that are not involved in Stl binding, including the β' jaw domain and non-conserved sequence insertions βSI1 in the lobe2 and β'SI3 in TL of Eco RNAP, and β'NCD1 in the clamp domain of Tth RNAP) are removed to make the Stl-binding pocket more visible.

**B**

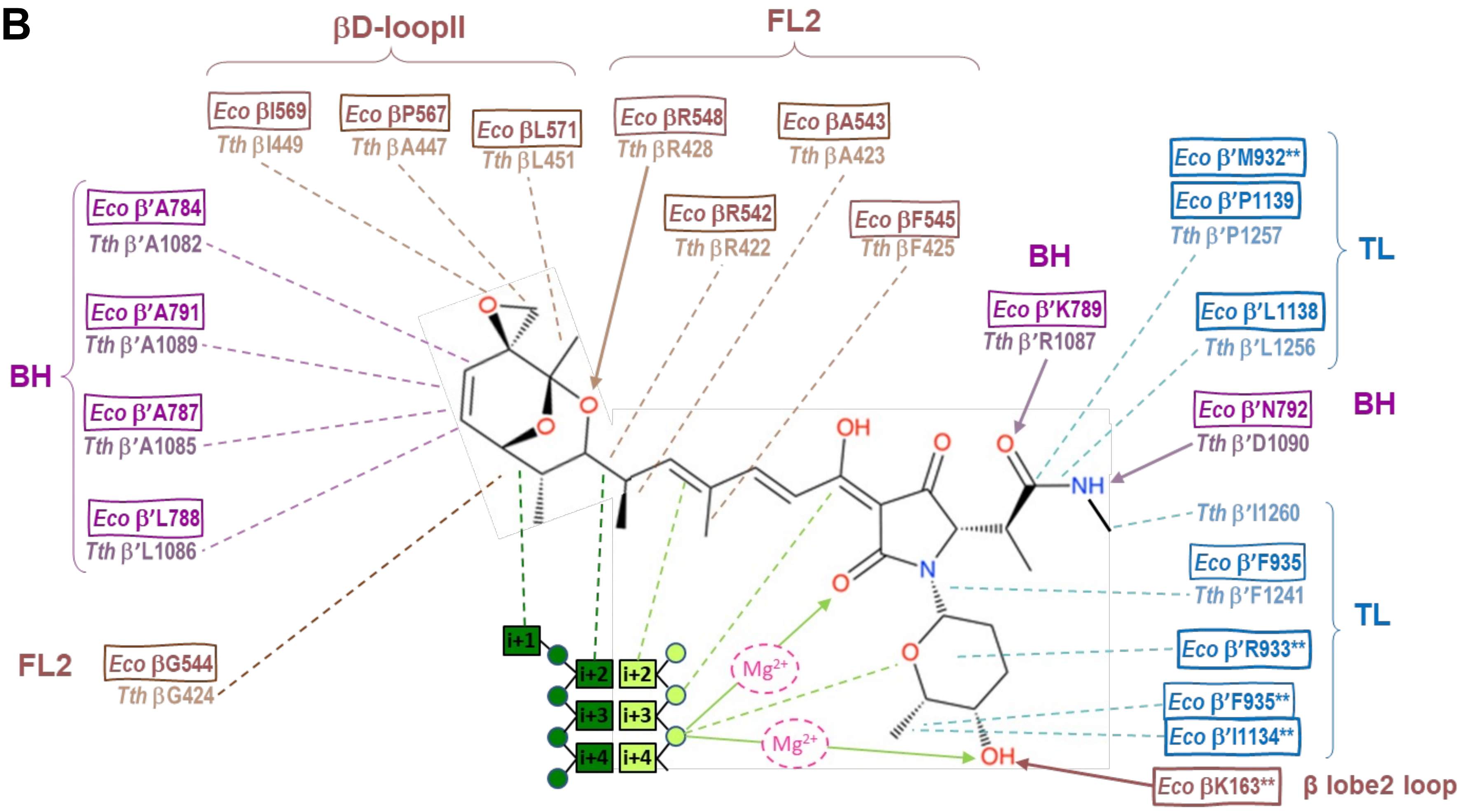


**Suppl. Figure S2. (B).** Schematic diagram showing amino acid residues and DNA nucleotides that contribute to the Stl-binding pocket in Tth EC-Stl and modeled Eco EC-Stl and Ecol EC-GreB-Stl complexes. The polar and van der Waals interactions are shown as solid arrows and dashed lines, respectively. Equivalent residues from Tth and Eco RNAPs that make similar contacts with Stl are grouped. Residues marked with asterisks indicate contact that Stl makes only in modeled EC-Stl-GreB complex. The residue color coding is the same as in (A). The Figure is modified from Suppl Fig. 8 of Vassylyev et al., 2007.

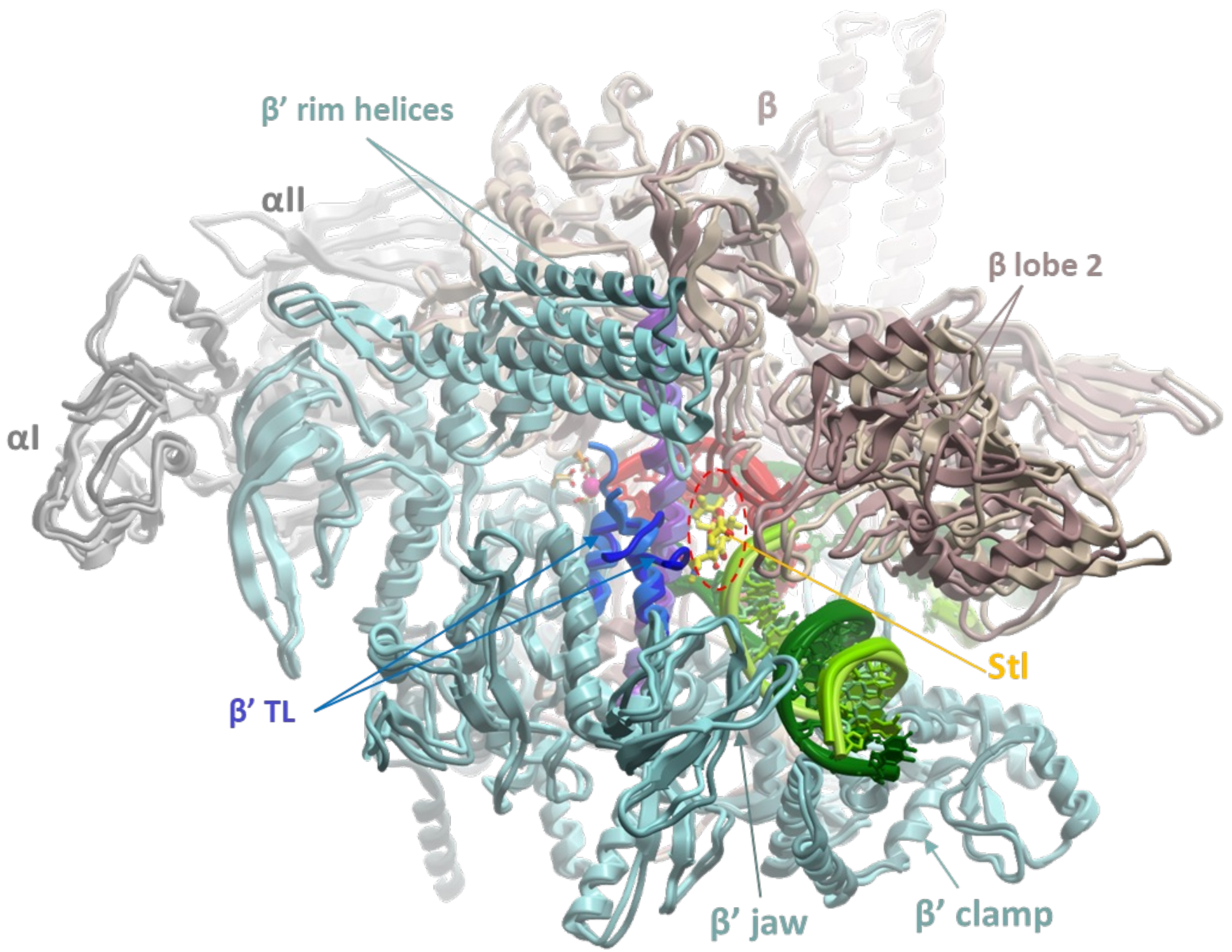


**Suppl. Figure S3.** Spatial rearrangements of Eco RNAP domains in EC induced by GreB, which may stabilize Stl binding. Structural alignment of Eco EC with the bound substrate, CTP (PDB: 6RH3, lighter colors), and Eco EC with bound GreB (PDB: 6RI7, darker colors) (Abdelkareem et al., 2019). The structures are shown as colored ribbons. The view and the coloring of the active center mobile elements are the same as in Suppl Figure S2. RNAP subunits and structural elements that undergo rearrangements upon GreB binding are indicated. GreB and the non-conserved part of the TL (β'SI3 domain) are not shown to make the tight Stl-binding pocket (indicated by a red dashed oval) more visible. The position of Stl in Eco EC is modeled based on the structural alignment of Eco EC and Tth EC-Stl complexes (see Suppl. Figure S2).

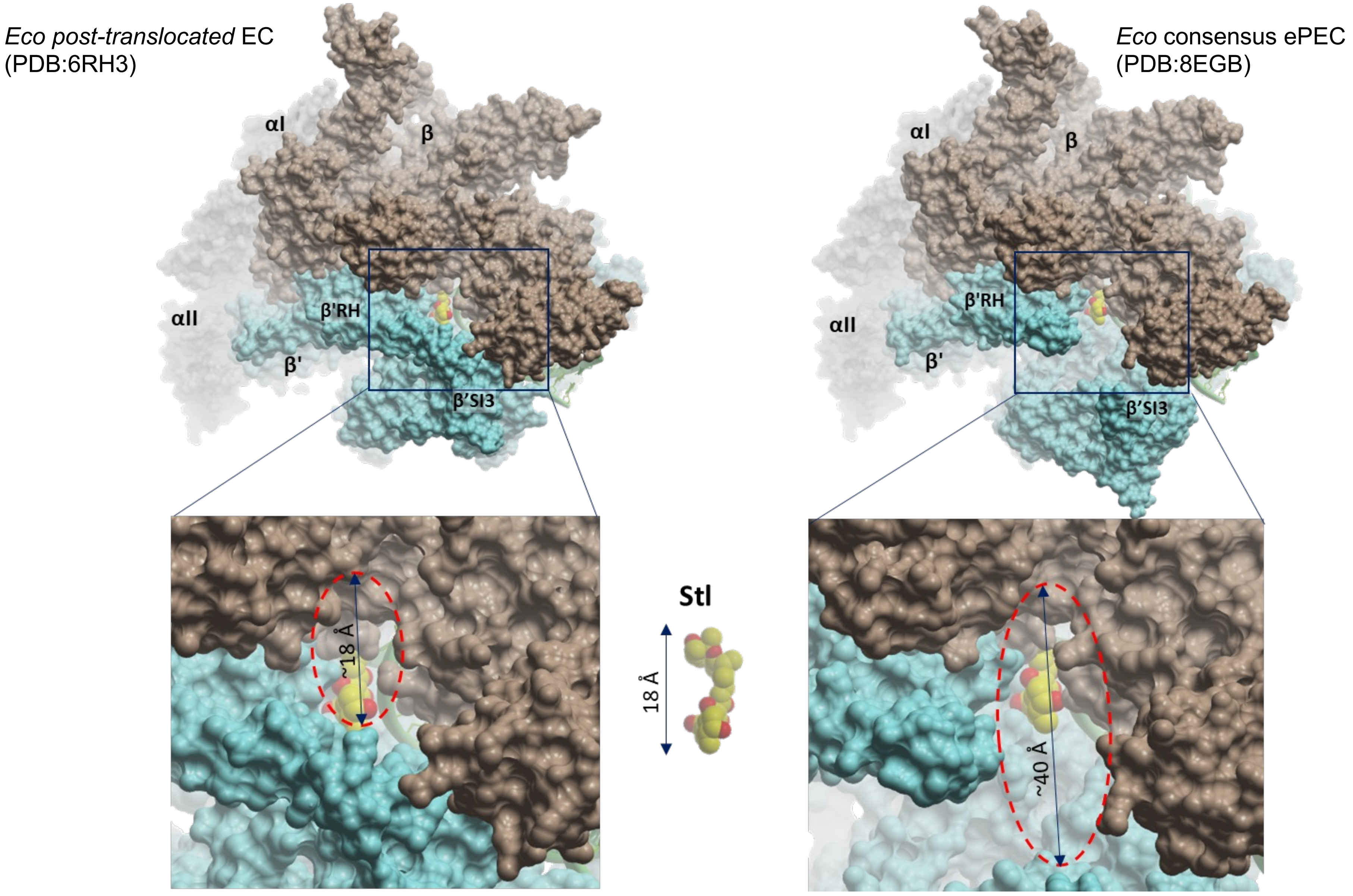


**Suppl. Figure S4.** Cryo-EM structures of post-translocated Eco EC (PDB: 6RH3) (Abdelkareem et al., 2019) and Eco consensus ePEC (PDB: 8EGB) (Kang et al., 2003) shown as solvent-accessible surfaces with superimposed Stl (CPK-colored balls). The surfaces of RNAP subunits are colored as in Fig. 4B. The spatial locations of the two mobile domains, β' rim helices (RH) and β'SI3, are indicated. The insets below show the magnified view of the Stl-entry channel in each structure: partially open (left panel) and fully open (right panel). Red dashed ovals with double-ended arrows indicate the approximate dimensions of the channel opening. The CPK structure of free Stl is shown between the two panels in the same scale.
